# Alpha-synuclein induces epigenomic dysregulation of glutamate signaling and locomotor pathways

**DOI:** 10.1101/2021.06.12.448150

**Authors:** Samantha L. Schaffner, Zinah Wassouf, Diana F. Lazaro, Mary Xylaki, Nicole Gladish, David T. S. Lin, Julia MacIsaac, Katia Ramadori, Julia M. Schulze-Hentrich, Tiago F. Outeiro, Michael S. Kobor

## Abstract

**Background:** Mutations and multiplications in the gene encoding for alpha-synuclein are associated with Parkinson’s disease (PD). However, not all individuals with alpha-synuclein variants develop PD, suggesting that additional factors are involved. We hypothesized that increased alpha-synuclein might alter epigenetic regulation of PD pathways.

**Objectives:** To identify genome-wide DNA methylation and hydroxymethylation changes induced by overexpression of two alpha-synuclein variants in human dopaminergic neurons, and to relate these to the corresponding transcriptome.

**Methods:** We assessed DNA methylation and hydroxymethylation at >850,000 CpGs using the EPIC BeadChip in LUHMES cells differentiated to dopaminergic neurons. Control LUHMES neurons, LUHMES neurons overexpressing wild type alpha-synuclein, and LUHMES neurons overexpressing A30P alpha-synuclein were compared. We used SMITE network analysis to identify functionally related genes with altered DNA methylation, DNA hydroxymethylation, and/or gene expression, incorporating LUHMES H3K4me1 ChIP-seq to delineate enhancers in addition to the default promoter and gene body regions.

**Results:** Using stringent statistical thresholds, we found that increased expression of wild type or A30P mutant alpha-synuclein induced DNA methylation changes at thousands of CpGs and DNA hydroxymethylation changes at hundreds of CpGs. Differentially methylated sites in both genotypes were enriched for several processes including movement-associated pathways and glutamate signaling. For glutamate and other signaling pathways (i.e. PDGF, insulin), this differential DNA methylation was also associated with transcriptional changes.

**Conclusions:** Our results indicated that alpha-synuclein altered the DNA methylome of dopaminergic neurons, influencing regulation of pathways involved in development, signaling, and metabolism. This supports a role for alpha-synuclein in the epigenetic etiology of PD.

## Introduction

Parkinson’s disease (PD) is characterized pathologically by the presence of alpha-synuclein (aSyn)-rich Lewy bodies in the brain. The function of aSyn is still unclear, as different cellular roles have been reported, including SNARE complex assembly, regulation of neuronal differentiation, glucose levels, and dopamine biosynthesis, as well as modulation of calmodulin activity/G-protein-coupled receptor kinase activity^1^. aSyn point mutations can impair one or several of these processes, disrupting neuronal health; however, the mechanism leading to PD may differ according to the variant. For example, the A30P mutation occurs in the membrane binding domain of aSyn, which may affect its ability to act as a presynaptic chaperone and interact with membrane-bound receptors^2^. Although A30P aSyn is less likely to aggregate than wild type (WT) aSyn, this mutation can still result in PD by loss of aSyn function^2^. Conversely, multiplications of the SNCA gene can lead to excess protein production, promoting aggregations and fibrils that impair synaptic function and can lead to neuronal death^3^. In familial PD patients, duplications and triplications of SNCA have been reported^4^.

Although a clearer picture of the relationship between aSyn dysfunction and PD is forming, much remains to be elucidated. For example, PD is variably penetrant even among individuals with pathogenic SNCA mutations, suggesting that additional genes and the environment likely play a role in etiology^4^. Since different aSyn mutations can lead to PD through different mechanisms, it is also important to understand which aspects of these variants are unique and which are shared. Building on this, our group and others have shown that expression of wild type or A30P mutant aSyn in human neurons can cause transcriptional dysregulation of cell survival and DNA repair genes, which may be mediated by altered histone H3 acetylation^5–6^. Additionally, altered DNA methylation (DNAm) patterns have been observed both at the SNCA gene itself and genome-wide in blood, saliva, and brain of PD patients^7–11^.Taken together, this implies that genome-wide transcriptional and epigenetic dysregulation may be involved in PD susceptibility and pathogenesis.

Epigenetic studies of PD provide the opportunity to assess genetic and environmental influences on disease risk concurrently^12–18^. DNAm, which refers to the attachment of a methyl group to DNA (on the 5’ carbon of cytosine, frequently in the context of CpG dinucleotides), is a well-studied epigenetic mark that changes during development and is influenced by genes and environment^12–15,19^.Although DNAm patterns can be unrelated to mRNA expression patterns or laid down as a consequence of gene expression, DNAm can in some cases also impact transcription by altering the ability for transcription factors to bind to genes, changing chromosomal interactions, and influencing splicing^12,20–23^.This can alter regulation of pathways involved in disease susceptibility^24–26^. Additionally, DNAm patterns can also be useful as biomarkers to indicate aging, diseased states, or exposures regardless of association with transcription^27–33^.

Although DNAm is influenced by a myriad of factors that differ between individuals, the existing literature on DNAm in PD has primarily taken a case-control approach^7–11^. Additionally, the role of DNA hydroxymethylation (DNAhm), which accounts for up to 40% of all modified cytosines in brain tissue, remains unclear^34^. DNAhm is an oxidized form of DNAm, and an intermediate of the DNA de-methylation process; however, studies have suggested that DNAhm is also a stable, independent mark in brain^35^. The stability of DNAhm and its ability to influence transcription (by eliminating DNAm-protein interactions, introducing DNAhm-protein interactions, altering chromatin state, and altering splicing) suggest that it could impact neurological disease etiology^36–37^. It is important to distinguish DNAhm from DNAm, since conventional bisulfite conversion techniques measure both DNAhm and DNAm in one compound signal, which may bias interpretations^38^. DNAhm is stable in post-mortem formalin-fixed tissue, and some initial studies have characterized DNAhm patterns in brains of deceased PD patients^39^. Bulk DNAhm levels are unchanged in the cortex, substantia nigra, and brainstem of PD patients, while DNAhm levels and expression of TET2, the enzyme which forms this mark, are elevated in neurons from prefrontal cortex of PD patients^40–41^. Importantly, DNAm and DNAhm patterns can be influenced by genetics, and genetic alterations at loci such as SNCA influence PD susceptibility, which suggests that incorporating a genetic lens into future epigenetic studies of PD will be important^42–44^. To date, only a few studies have assessed the impact of genetic background on DNAm patterns in PD^45–47^.

One of the major challenges in human epigenetic studies of PD is accounting for the range of genetic, environmental, and lifestyle factors that can influence DNAm and DNAhm^42,48^. Furthermore, using postmortem tissue to study PD also typically presents the limitation of advanced disease. Model systems such as rodents or cell culture help to address these issues, providing a context where both genetics and environment can be tightly controlled, and the opportunity to examine early impacts of neurological disease in brain.

In this study, we characterized the influence of two molecularly distinct aSyn variants on the DNA methylome and hydroxymethylome of human dopaminergic neurons. We profiled genome-wide DNAm and DNAhm patterns in three groups of Lund human mesencephalic (LUHMES) cells differentiated into dopaminergic neurons: control LUHMES, LUHMES overexpressing WT aSyn at levels similar to those seen in SNCA multiplication carriers, and LUHMES overexpressing A30P aSyn. We additionally integrated our DNAm and DNAhm data with transcriptomic data from the same cells, incorporating H3K4me1 ChIP-seq to score DNA(h)m changes across gene regulatory features^6^. WT and A30P aSyn expression were associated with thousands of DNA(h)m changes, particularly affecting genes that regulate locomotory and glutamate signaling pathways. This suggested that familial PD-associated aSyn mutations have widespread epigenomic effects, and may contribute to molecular and transcriptional dysregulation associated with PD etiology and heterogeneity.

## Materials and Methods

### Generation of aSyn expressing LUHMES cells

LUHMES cells were a gift from Prof. Marcel Leist and were cultured and differentiated as previously described^6^ (Supplementary Methods). In brief, full-length human WT or A30P aSyn cDNA was cloned into a biscistronic lentiviral vector under a chicken beta-actin promoter, and transiently transfected into 293T cells. Proliferating LUHMES were infected with equimolar amounts of IRES-GFP, WT aSyn-IRES-GFP or A30P aSyn-IRES-GFP lentivirus. Infected cells were selected by FACS, and differentiated for eight days^6^(Supplementary Methods).

### EPIC BeadChip data generation and bioinformatic analysis

DNA was extracted from cell pellets (see Supplementary Methods), and 750 ng per sample of bisulfite (BS)- or oxidative bisulfite (oxBS)-converted DNA was run on Infinium HumanMethylationEPIC (EPIC) BeadChips (Illumina Inc.) according to the manufacturer’s instructions, producing data for 853,307 CpG and 2,880 CNG sites.

Beta values were generated from raw intensity signals using GenomeStudio software (Illumina) and exported into R 3.6 for data analysis. SNP control probes, cross-hybridizing probes, and low-quality probes were removed, leaving 813,589 EPIC probes in the final dataset^49–50^. oxBS and BS data were normalized separately with dasen, and batch effects were removed with ComBat^51^. Hydroxymethylation (hmC) values were calculated by subtraction, and a detectability threshold of 3.6% was set using the 95% quantile of resulting negative values ^52^. See Supplementary Methods for more detailed data processing information.

Differential methylation between control (n=7), WT aSyn (n=8), and A30P aSyn (n=8) cells was calculated by linear regression using limma with the model DNA(h)m ~ Genotype^53^. Individual sites were considered significant at FDR <= 0.05 and |delta beta| >= 0.05 (calculated by subtracting the mean beta value across all replicates of the group in question from the reference group, e.g. WT aSyn beta - Control beta). The skewness.norm.test function from the normtest R package was used to assess delta beta skewness, with 1000 Monte Carlo simulations^54^.

### Pyrosequencing

See Supplementary Methods.

### Gene ontology enrichment

Genes were assigned to CpGs if they belonged to the longest UCSC RefGene transcript annotated to each site, resulting in 30,325 unique coding- and non-coding gene annotations for the DNAm dataset and 23,190 unique gene annotations for the DNAhm dataset. Gene ontology enrichment was performed with the over-representation (ORA) method in ermineR 1.0.1.9, using differentially methylated genes identified with the effect size and significance thresholds previously mentioned as input^55^. Genes that were differentially methylated or hydroxymethylated in both WT aSyn and A30P aSyn cells in either direction were removed from the input list and analyzed separately.

### Chromatin immunoprecipitation sequencing (ChIP-seq) and peak calling

See Supplementary Methods.

### Multi-omic integration

DNAm, DNAhm, and RNA-seq data were integrated using SMITE in R 3.6, with adjusted p-values as significance input, delta betas as effect size input for DNAm and DNAhm, and log2FC values for RNA-seq^56^(Supplementary Methods). H3K4me1 ChIP-seq peaks were used to define enhancers. The following weights were applied during scoring: expression, 0.4; enhancer DNAm, 0.125; promoter DNAm, 0.125; body DNAm, 0.1; enhancer DNAhm, 0.125; promoter DNAhm, 0.125; body DNAhm, 0.1. Gene scores were annotated to a REACTOME protein-protein interaction network.

## Results

### aSyn overexpression altered genome-wide DNAm patterns in dopaminergic neurons

In order to evaluate the epigenome-wide impacts of expressing aSyn in dopaminergic neurons, we assessed DNAm differences between control, WT aSyn, and A30P aSyn cells at 813,589 EPIC probes. We selected an effect size threshold of |delta beta| >= 0.05 to be well above technical noise (maximum 2.2% RMSE between technical replicates). Firstly, we sought to confirm whether overexpression of aSyn was associated with reduced DNAm at the first intron of SNCA, as previously reported in PD patients^10–11^; this was true for both aSyn genotypes (Fig. 1). We next assessed genome-wide DNAm changes induced by overexpression of each aSyn variant (Fig. 2). When WT aSyn cells were compared to control, 18,521 sites had decreased DNAm and 10,812 sites had increased DNAm (*padj* <= 0.05, Fig. 2A, Table S2). In A30P aSyn cells, 3,091 probes had decreased DNAm compared to control while 3,438 probes had increased DNAm (Fig. 2A, Table S3). When compared to WT aSyn cells, A30P aSyn cells had 3,014 sites with decreased DNAm and 6,666 sites with increased DNAm (Fig. 2A, Table S4). DNAm changes in all comparisons were significantly skewed to one direction, and the bias in effect direction was seen across all genomic features (*p* < 2.2e-16, Table S5). To confirm an example from our findings by pyrosequencing, we selected a region of the *TUBA8* gene that included the top two probes with the largest change in DNAm across both genotypes; differential DNAm at these two positions, one additional position on the array, and four positions not on the array were all confirmed (Fig. S1, Table S8).

**Figure 1.**
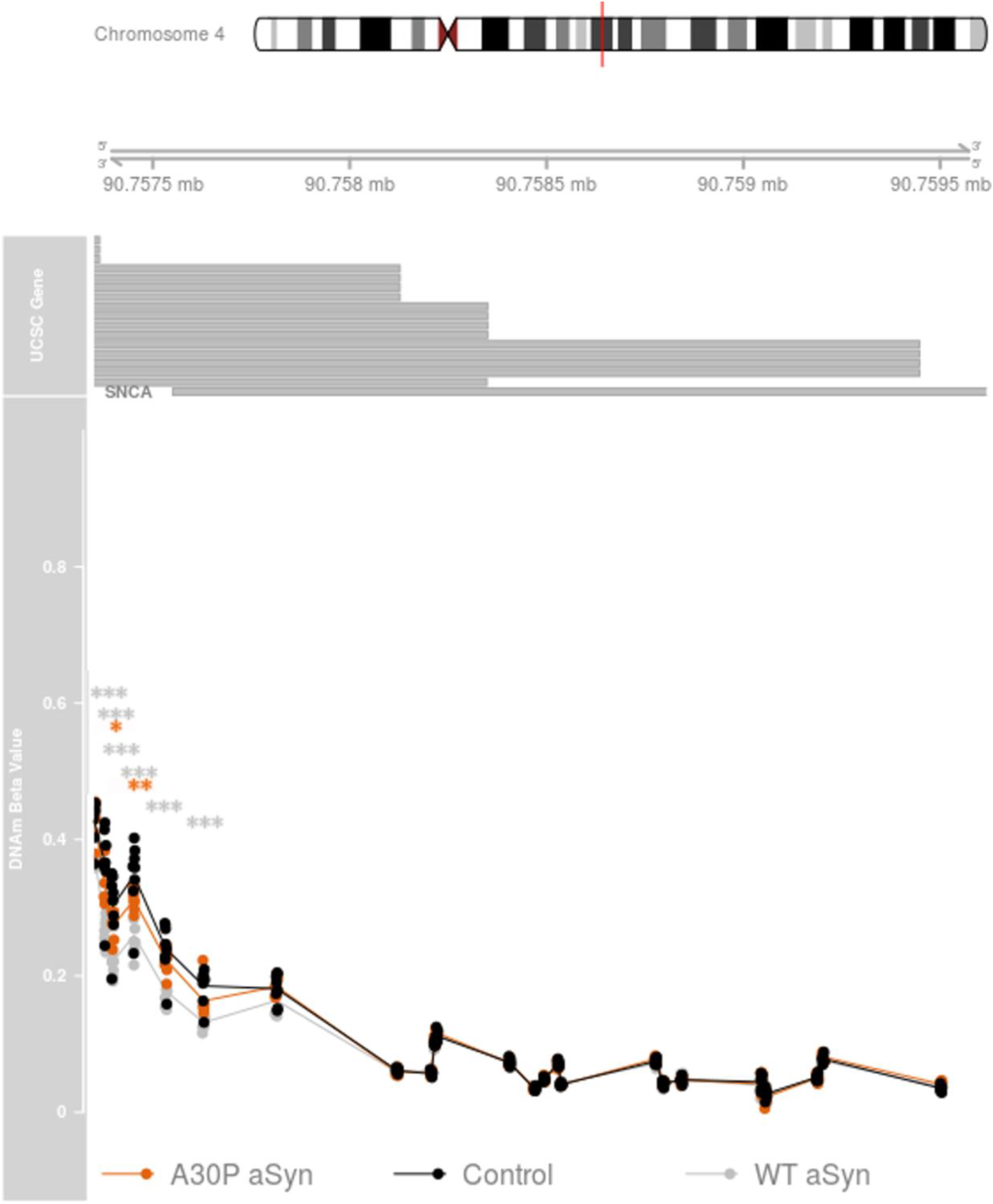
aSyn overexpression was associated with decreased DNAm at the SNCA 5’ UTR. Top: UCSC hg19 coordinates are shown; grey boxes represent *SNCA* transcripts. Bottom: DNAm beta values are shown for each sample, coloured by genotype. Black: control cells; pale grey: WT aSyn cells; orange: A30P aSyn cells. * *padj* < 0.05, ** *padj* < 0.005, *** *padj* < 0.001.

**Figure 2.**
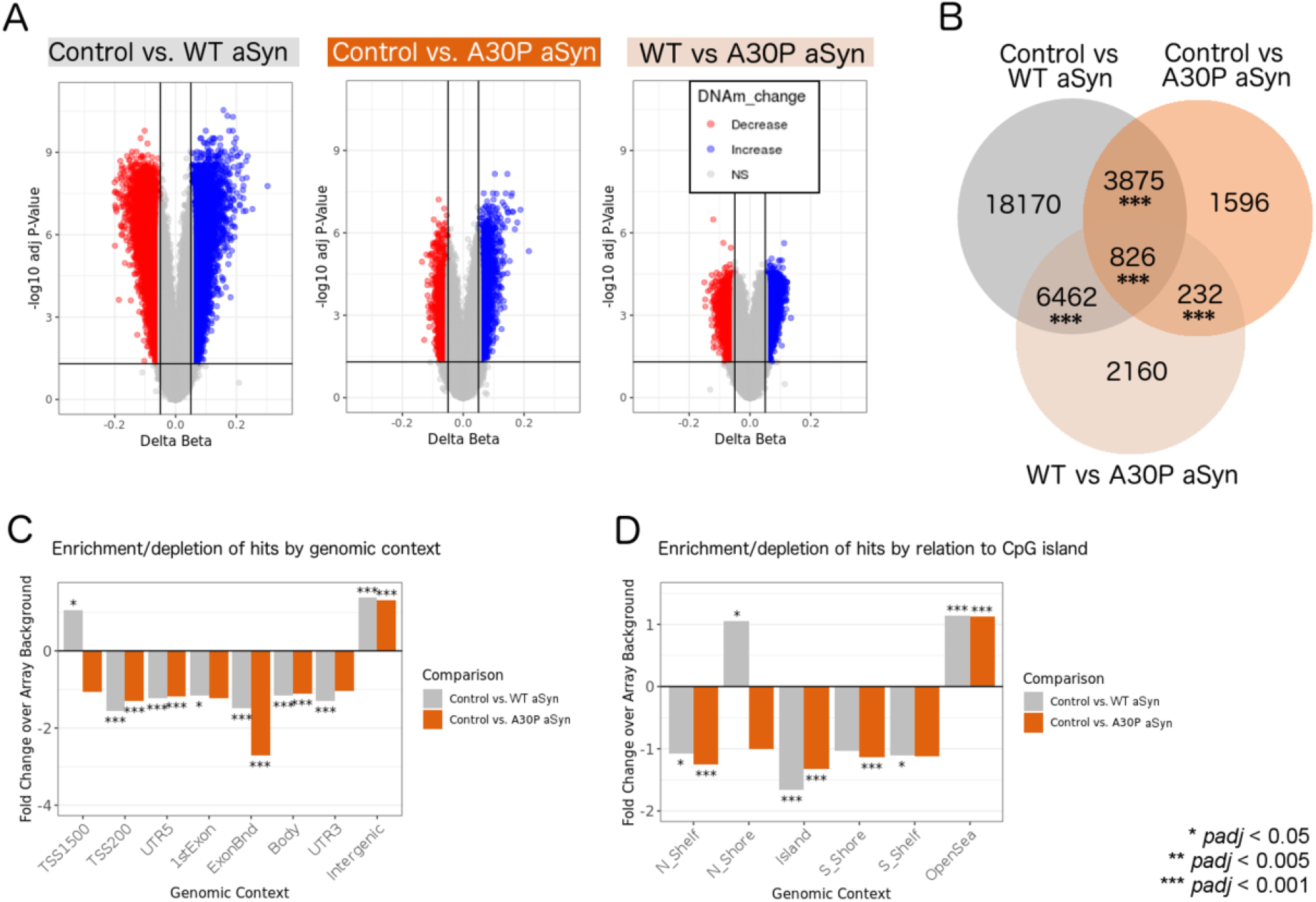
Overexpression of WT and A30P mutant aSyn altered the DNA methylome of dopaminergic neurons. (A) Volcano plots comparing DNAm patterns between control and WT aSyn LUHMES, control and A30P aSyn LUHMES, and WT aSyn and A30P aSyn LUHMES. Red: sites with decreased DNAm; blue: sites with increased DNAm (|delta beta| >= 0.05 and *padj* <= 0.05). (B) Number of differentially methylated probes unique to each comparison and probes shared between comparisons. * *padj* < 0.05, ** *padj* < 0.005, *** *padj* < 0.001 (10,000 permutations). (C) Relative enrichment/depletion of differentially methylated sites across genomic contexts, permuted against array background. Grey: control vs. WT aSyn analysis; orange: control vs. A30P aSyn analysis. * *padj* < 0.05, ** *padj* < 0.005, *** *padj* < 0.001 (10,000 permutations). (D) Relative enrichment/depletion of differentially methylated sites by relation to CpG island, permuted against array background. Grey: control vs. WT aSyn analysis; orange: control vs. A30P aSyn analysis.

The proportion of differentially methylated probes shared across all comparisons and any two of three comparisons was larger than expected by chance, and in most cases, WT aSyn cells had more pronounced differences from controls than A30P aSyn cells (Fig. 2B, *padj* < 0.005, 10,000 permutations; Fig. S2). Interestingly, WT and A30P aSyn overexpression primarily affected CpGs outside of genes; however, trends for differentially methylated probe feature enrichment differed between genotypes for TSS1500 regions and CpG North Shores (Fig 2C-D, both enriched in WT cells and depleted in A30P cells).

### aSyn overexpression altered genome-wide DNAhm patterns to a lesser degree than DNAm patterns

After assessing genome-wide DNAm alterations, we examined differential DNAhm at 233,440 sites that passed our detection threshold of 3.6% (calculated according to the 95% quantile of negative hmC values after subtraction; see Methods). Fewer DNAhm changes for each comparison were observed as compared to DNAm, most of which were increases; these increases in DNAhm were significant when tested for skewness (*p* < 2.2e-16, Fig. 3, Fig. S3, Tables S9-S11). The overlap of 16 CpG sites between control vs. WT aSyn and control vs. A30P aSyn was not higher than expected by chance (Fig. 3B, *padj* > 0.05, 10,000 permutations).

**Figure 3.**
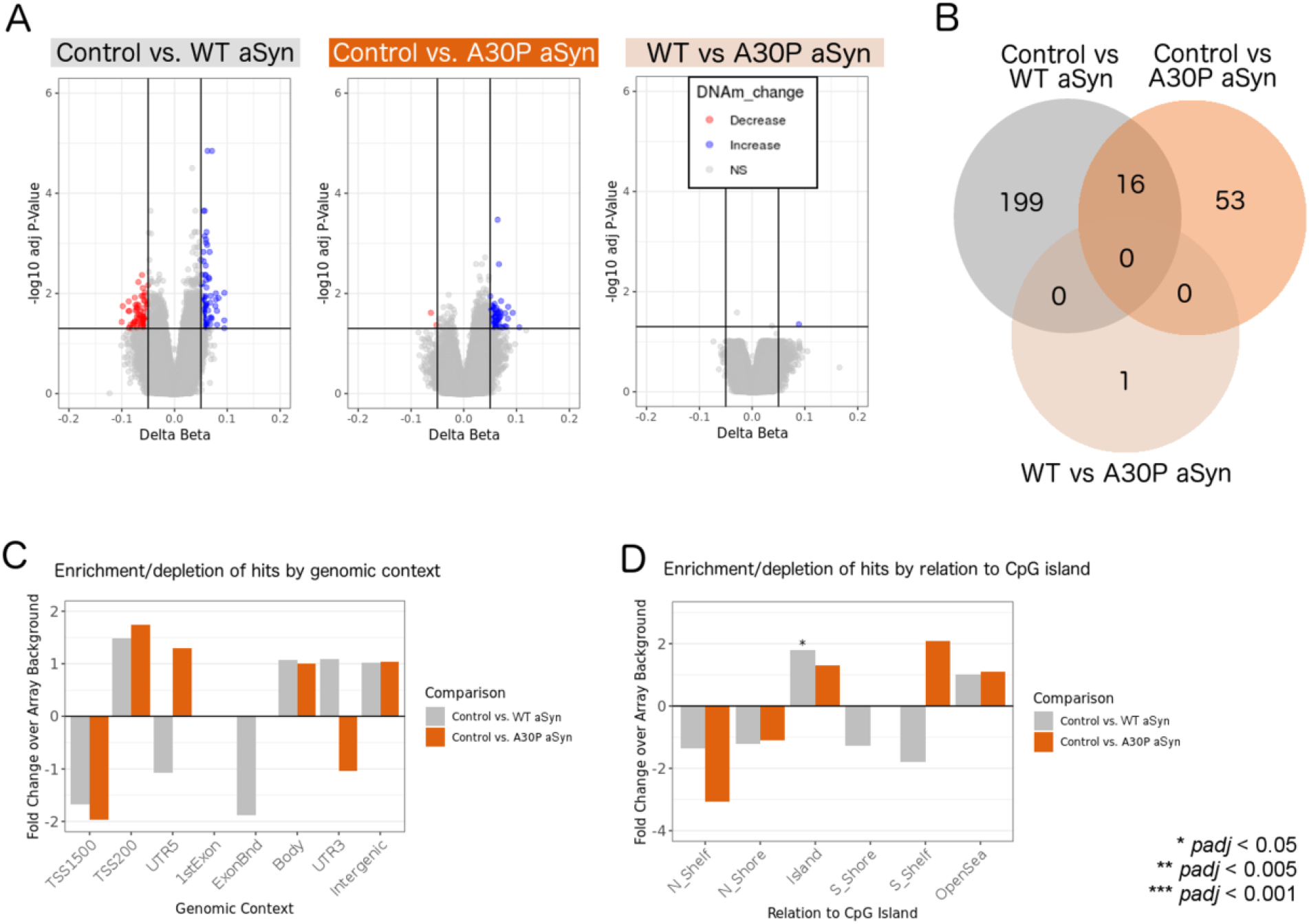
WT and A30P aSyn overexpression were correlated with increased DNA hydroxymethylation levels in dopaminergic neurons. (A) Volcano plots comparing DNAhm patterns between control and WT aSyn LUHMES, control and A30P aSyn LUHMES, and WT aSyn and A30P aSyn LUHMES. Red: sites with decreased DNAm; blue: sites with increased DNAm (|delta beta| >= 0.05 and *padj* <= 0.05). (B) Number of differentially hydroxymethylated probes unique to each comparison and probes shared between comparisons. All *padj* > 0.05 (10,000 permutations). (C) Relative enrichment/depletion of differentially hydroxymethylated sites across genomic contexts, permuted against array background. Grey: control vs. WT aSyn analysis; orange: control vs. A30P aSyn analysis. (D) Relative enrichment/depletion of differentially hydroxymethylated sites by relation to CpG island, permuted against array background. Grey: control vs. WT aSyn analysis; orange: control vs. A30P aSyn analysis. * *padj* < 0.05 (10,000 permutations).

### WT and A30P aSyn altered DNAm at movement-associated and glutamate signaling pathway genes

To identify possible functional consequences associated with the DNAm and DNAhm changes in each genotype, we performed gene ontology enrichment analysis on all differentially methylated or hydroxymethylated genes within each comparison using over-representation analysis in ermineR^57^. 34 GO biological processes were enriched in differentially methylated sites shared between the control vs. WT aSyn and control vs. A30P aSyn comparisons (Fig. 4A, *padj* < 0.05). 442 CpG sites were annotated to the top GO term, “locomotory behavior,” and differentially methylated in at least one genotype (420 sites in WT aSyn cells and 89 sites in A30P aSyn cells, Fig. 4B-E). 129 differentially methylated sites were annotated to the second-highest ranked GO term, “glutamate receptor signaling pathway” (127 sites in WT aSyn cells and 18 sites in A30P aSyn cells, Fig. S5). No significant multifunctionality-corrected enrichments were observed for differentially methylated or hydroxymethylated sites in any of the other groups.

**Figure 4.**
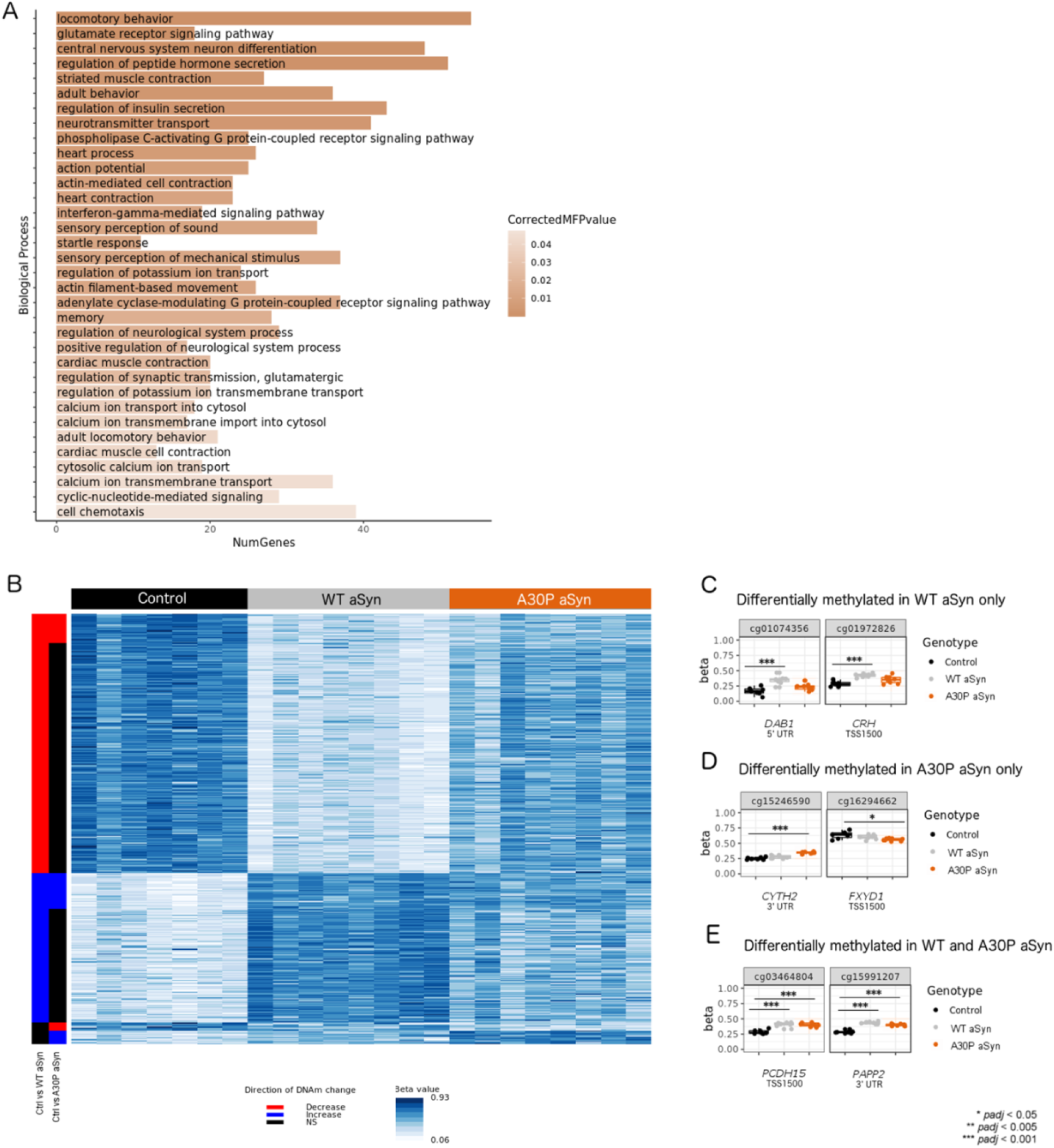
Differentially methylated genes in WT and A30P aSyn cells were enriched for locomotory behavior and glutamate receptor signaling functions. (A) GO biological process terms enriched at multifunctionality-corrected, multiple test-corrected *p*-value (CorrectedMFPvalue) < 0.05 in genes shared between control vs. WT aSyn and control vs. A30P aSyn DNAm analyses. (B) Heat map showing beta values for all probes annotated to the “locomotory behavior” pathway and differentially methylated in at least one comparison. Row labels, left to right: Probes that passed significance thresholds (|delta beta| >= 0.05 and *padj* <= 0.05) in control vs WT aSyn comparison (decreased DNAm in WT aSyn: red; increased DNAm in WT aSyn: blue; non-significant: black). Probes that passed significance thresholds in control vs A30P aSyn comparison (decreased DNAm in A30P aSyn: red; increased DNAm in A30P aSyn: blue; non-significant: black). (C) Representative examples of locomotory behavior-related CpG sites differentially methylated only in WT aSyn cells. (D) Locomotory behavior-related CpG sites differentially methylated only in A30P aSyn cells. (E) Examples of locomotory behavior-related CpG sites differentially methylated in both genotypes. Black: control cells; grey: WT aSyn cells; orange: A30P aSyn cells. * *padj* < 0.05, ** *padj* < 0.005, *** *padj* < 0.001.

### aSyn impacted epigenetic and transcriptional regulation of glutamate, NOTCH, insulin, PDGF, and SHH signaling network genes

We next queried genes and pathways that associated with changes in both the epigenome and the transcriptome in WT and A30P aSyn cells. These loci represent candidates where DNAm and/or DNAhm may be regulating gene expression, and it is important to identify them in order to understand the molecular etiology of PD. Firstly, we compared the DNAm and DNAhm hits found in this study against previously identified differentially expressed genes^6^(863 genes in WT aSyn cells and 1,315 genes in A30P aSyn cells, |log2FC| > 0.5 and *padj* < 0.01). Out of 10,798 genes with data for all three modifications in WT aSyn cells, seven displayed differential DNAm, DNAhm, and expression, including an ionotropic glutamate receptor (*GRIK2*; Table S15). In A30P cells, two out of 10,831 genes had concurrent changes to these modifications (*TSPEAR* and *SGPP2*, Table S16). The relationship between DNAm, DNAhm, and expression varied depending on the gene and the CpG site in question (Tables S15-S16).

We additionally employed SMITE, which captures modules of functionally related genes with changes to at least one of either DNAm, DNAhm, or gene expression in each comparison^56^. We included H3K4me1 ChIP-seq data from the same LUHMES in our SMITE workflow, allowing us to consider DNAm and DNAhm changes at enhancers (in addition to default promoter and gene body regions), and weighted the importance of each modification (expression: 0.4, DNAm: 0.35, DNAhm: 0.25). In WT aSyn cells, 18 modules were identified (Table S17, Fig. S7). In A30P aSyn cells, 24 modules were identified (Table S18, Fig. S8).

Glutamate receptor signaling modules discovered using SMITE stood out with more concurrent changes to DNAm, DNAhm, and transcription in both genotypes. Two modules centered around *GRIK2* were each significantly altered in WT and A30P cells, matching with the direct comparison of differentially (hydroxy)methylated versus differentially expressed genes. There was a negative correlation between *GRIK2* promoter DNAm and mRNA levels in both genotypes (Fig. 5).

**Figure 5.**
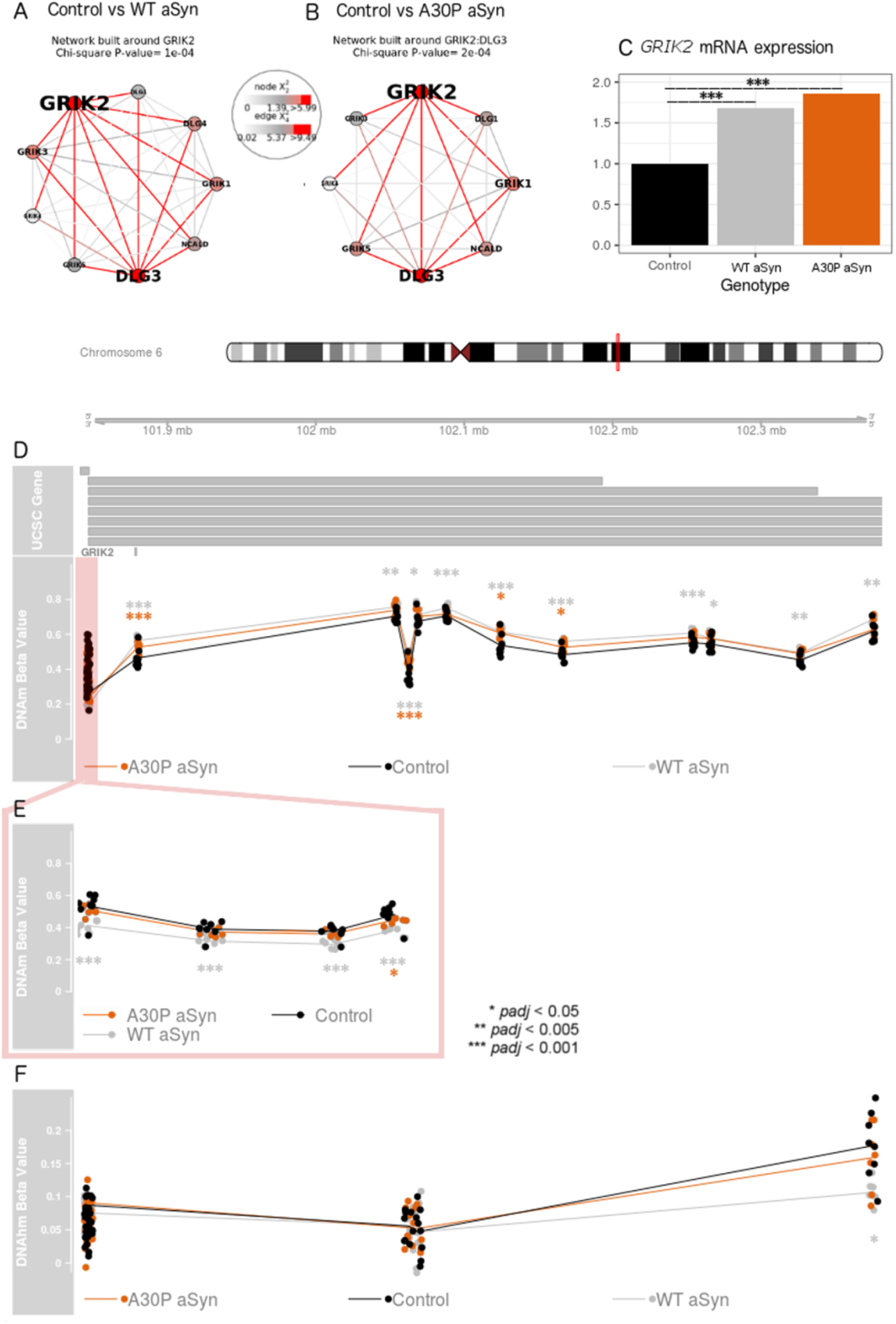
*GRIK2* was differentially methylated, hydroxymethylated, and expressed in LUHMES overexpressing either WT or A30P aSyn. (A) GRIK2 protein-protein interaction network which showed differential DNAm, DNAhm, and expression in control vs. WT aSyn cells for the underlying genes. (B) GRIK2:DLG3 protein-protein interaction network which showed differential DNAm, DNAhm, and expression in control vs. A30P cells for the underlying genes. (C) Relative expression of *GRIK2* mRNA in each genotype, normalized to control cells. (D) Top: UCSC hg19 coordinates are shown, with *GRIK2* transcripts below. Bottom: DNAm beta values are shown for each sample, coloured by genotype. Black: control; grey: WT aSyn; orange: A30P aSyn. (E) DNAm at the *GRIK2* TSS200 region. (F) DNAhm levels for EPIC array probes across the *GRIK2* gene. * *padj* < 0.05, ** *padj* < 0.01, *** *padj* < 0.005.

## Discussion

Elucidating the mechanisms by which PD develops and progresses in different individuals is paramount to developing preventative strategies and treatments. Here, we explored whether DNAm and DNAhm contributed to interindividual differences in PD etiology among carriers of aSyn variants. We assessed the impact of overexpressing WT or A30P aSyn on genome-wide DNAm and DNAhm patterns in dopaminergic neurons, the primary cell type affected in PD. We found that overexpressing either aSyn variant induced thousands of DNAm changes and hundreds of DNAhm changes in pathways related to PD and neurodegeneration, and that WT aSyn particularly impacted glutamate receptor signaling genes on the epigenetic and transcriptional level. Distinct characteristics of each aSyn protein may explain why both similar and unique effects on DNA(h)m were observed. This study enhances our understanding of the wide-ranging genomic impacts of different aSyn forms and illuminates further possible molecular mechanisms for PD.

The dysregulation of locomotor behaviour pathway genes was reassuring and indicative of the relevance and validity of our approach since this is one of the major systems affected in PD. Excessive glutamatergic transmission can also play a role in PD, causing excitotoxicity in dopaminergic neurons^58^. Preclinical studies have implicated glutamate receptors as therapeutic targets for PD, and regulation of glutamate signaling genes may be influenced by lifestyle factors such as dietary exposure to neurotoxins or tea polyphenols^16,59–61^. Our results thus provide evidence from an epigenetic perspective to expand the existing research on the role of glutamate signaling in PD etiology and prevention, and add a role for aSyn in this pathway.

Aside from these core similarities in the pathways affected by each aSyn variant, we also saw unique effects of WT and A30P aSyn on the DNA methylome and hydroxymethylome. Broadly, many more DNAm and DNAhm changes were seen in WT aSyn cells as compared with A30P cells. This was surprising in light of our previous work, which identified that A30P cells are affected to a greater extent on the transcriptional level^6^. It was also unexpected that almost no changes to non-CpG methylation were observed with aSyn overexpression (one differentially methylated CH probe in WT vs A30P aSyn cells, data not shown) despite known enrichment of non-CpG methylation in neurons^62–63^. Several mechanisms might explain this. Firstly, DNA damage occurs to a greater extent in WT aSyn LUHMES neurons than in A30P aSyn LUHMES neurons, and it is possible that some DNAm changes in WT cells are a reflection of this damage and/or subsequent repair attempts^6,64^. Following DNA damage, protective mechanisms in areas of active transcription maintain DNA repair and restore epigenetic modifications; this may also explain why we observed more DNAm changes in intergenic regions and less in intragenic regions than expected by chance in both genotypes^64–65^.

Differences in characteristics of each protein could also explain the higher number of DNAm alterations in WT cells. WT aSyn could influence DNAm through various means, such as binding to membrane-bound G-coupled protein receptors and initiating signaling cascades; facilitating SNARE complex assembly at presynaptic membranes, which could allow neurotransmitters to activate signaling cascades at the postsynaptic neuron; and binding to DNA^3,66^. The A30P mutation prevents aSyn from binding to membranes, removing some of these avenues^4^.

WT aSyn can sequester Dnmt1 from the nucleus in mice, resulting in global loss of DNA methylation in the brain^67^. If there is a difference in DNMT1 sequestration between aSyn variants, this could reduce the capacity for maintenance of DNAm in a non-specific manner. aSyn aggregation - which is more likely with the WT protein - has also been demonstrated to increase this sequestration^68^. Although A30P aSyn is more likely to localize to the nucleus than WT aSyn, the ability of A30P protein to sequester Dnmt1 has not been assessed^66,68–69^. In addition to influencing the DNAm loss in WT cells and gain in A30P cells, this potential difference in nuclear aSyn could also affect the amount of DNAm at transcription-associated regions, due to its ability to bind DNA^66,68–69^.

In contrast to DNAm, patterns of differential hydroxymethylation were similar between aSyn genotypes, and fewer DNAhm changes were seen at our statistical thresholds. This was expected due to the fetal origin of the LUHMES cell line^36,70^. The opposite direction of change seen for DNAm and DNAhm at the same loci was also expected, since DNAm must be oxidized in order for DNAhm to form^71^. We are unable to determine from our study whether this DNAhm was a transient intermediate or represents a stable epigenetic mark; however, correlating DNAhm patterns with gene expression levels provided some insight on loci where DNAhm was associated with transcription^35^.

It is important to understand the interplay between DNAm, DNAhm, and gene expression to gain insight into the molecular etiology of PD. Changes to DNAm may be unrelated to transcription, or precede or follow changes to transcription^12,20–21^. In the former case, DNAm may represent a target for modulation of gene expression in order to prevent, for example, upregulation of glutamate signaling genes in individuals with aSyn multiplications. In the latter case, DNAm is more likely to represent a biomarker of expression. The relatively low correlation between DNA(h)m changes and expression changes in this study (<0.1% of genes in either genotype) agrees with previous literature, which has reported correlations with expression at only 0.6-8% of CpG sites in blood and up to 0.3% of CpG sites in brain^72–76^. However, the few loci that did have transcriptional changes correlated with DNA(h)m changes may still represent either functionally relevant targets for PD or potential biomarkers for neurodegeneration-relevant pathways.

Although this study provided insights into the epigenomic impact of aSyn overexpression, several limitations should be considered. The *SNCA* transgene integrated at random positions in the genome, which could affect DNAm at a regional level. We used eight replicates of each LUHMES genotype and seven replicates of control LUHMES for EPIC array profiling, with each replicate consisting of a pool of thousands of cells, in order to represent a range of integration positions. Culturing cells can also influence DNAm^77–79^. Our LUHMES replicates were obtained from a range of passages and culture dishes, and the effect of passage was regressed out before differential DNAm/DNAhm analysis. This reduced bias due to passage, but may not have completely eliminated it. Additionally, determining DNAhm using a combination of oxidative bisulfite- and bisulfite-converted samples introduced technical noise, which led us to reduce the number of sites for DNAhm analysis^52^. Finally, the LUHMES cells used in this study may not be fully representative of dopaminergic neurons in the brains of PD patients due to their fetal origin, differentiation in tissue culture, and construct-based overexpression of both aSyn forms as opposed to genetic multiplication or A30P point mutation.

Overall, we demonstrated that WT and A30P aSyn overexpression each had significant impacts on the DNA methylome of human dopaminergic neurons, and that only a small proportion of these DNAm changes occur concurrently with DNAhm and gene expression changes to neurodegeneration-related genes. This study contributes to our understanding of the molecular genetic etiology of PD and lays groundwork for future studies to investigate whether epigenetic and transcriptional changes associated with increased expression or mutations in aSyn may be reversible with environment and/or lifestyle factors.

## Author Roles

1. Research project: A. Conception, B. Organization, C. Execution; 2. Statistical Analysis: A. Design, B. Execution, C. Review and Critique; 3. Manuscript: A. Writing of the first draft, B. Review and Critique.

S.L.S: 1A, 1B, 1C, 2A, 2B, 3A.

Z.W.: 1B, 1C, 2A, 2B, 3B.

D.F.L.: 1B, 1C, 3B.

M.X.: 1B, 1C, 3B.

N.G.: 2C, 3B.

D.T.S.L: 1C, 3B.

J.M.: 1C, 3B.

K.R.: 1C, 3B.

J.M.S.H.: 1A, 1B, 3B.

T.F.O: 1A, 1B, 3B.

M.S.K.: 1A, 1B, 2C, 3B.

