## Supplementary File 1 for "Alpha-synuclein induces epigenomic dysregulation of glutamate signaling and locomotor pathways"

**Supplementary Methods**

**LUHMES culturing and differentiation**

Proliferating cells were cultured in Advanced Dulbecco’s modified Eagle’s medium/F12 (DMEM/F12, Gibco) supplemented with N2 (Gibco), 2 mM L-glutamine (Gibco) and 40 ng/mL recombinant basic fibroblast growth factor (bFGF, R&D Systems), in pre-coated flasks (Corning) with 50 μg/mL PLO (Sigma-Aldrich), at 37ºC in a humidified 5% CO2 atmosphere.

Differentiation was achieved by replacing the proliferating medium with DMEM/F12 supplemented with N2 (Gibco), 2 mM L-glutamine (Gibco), 1 μg/mL tetracyclin (Sigma-Aldrich), 1 mM dibutyryl cAMP (cAMP, Sigma Aldrich) and 2 ng/mL recombinant human GDNF (R&D Systems). After two days in culture (pre-differentiation), cells were trypsinized and seeded into plates pre-coated with 50 μg/mL PLO and 1 μg/mL fibronectin (Sigma-Aldrich). On the fifth day of differentiation one third of culture medium was refreshed. Cells were harvested on the eighth day of differentiation.

**Chromatin immunoprecipitation sequencing (ChIP-seq)**

Chromatin and DNA were cross-linked by adding formaldehyde to the culture dish to a final concentration of 1% and incubating for 10 min at 25°C. The cross-linking reaction was quenched with glycine to a final concentration of 0.125 M at 25°C. Cells were then collected, washed with cold PBS and pelleted by centrifugation at 900 rcf for 5 min at 4°C. Cell pellets were resuspended in RIPA SDS buffer (140 mM NaCl, 1 mM EDTA at pH 8.0, 1% TritonX-100, 0.1% sodium deoxycholate, 10 mM Tris-Cl at pH 8.0, 1% SDS) supplemented with Roche complete protease inhibitors, incubated at 4°C for 10 min, and then used for shearing in Covaris® instrument. The sheared chromatin was cleared by centrifugation at 16 000 rcf for 5 min and an aliquot was used to check DNA size on an Agilent 2100 Bioanalyzer (Agilent Technologies). We proceeded with immunoprecipitation when DNA was sheared to fragments of 150-300 bp.

For chromatin immunoprecipitation, samples were diluted 10x in IP buffer (150 mM NaCl, 1% NP-40, 0.5% sodium deoxycholate, 50 mM Tris-HCl at pH 8.0, 2 mM EDTA at pH 8.0) supplemented with Roche complete protease inhibitors, pre-cleared with protein A magnetic beads (Dynabeads, Invitrogen) for 1h at 4°C, and then used for immunoprecipitation by histone modification-specific ChIP-grade antibody (anti H3 mono methyl K4 antibody, ab8895, Abcam). After overnight incubation at 4°C, protein A magnetic beads were added and incubated for 2 hours at 4°C. Beads were then washed twice with cold IP buffer with 0.1% SDS and protease inhibitors, 3x with wash buffer (100 mM Tris-HCl at pH 8.0, 500 mM LiCl, 1% v/v NP-40, 1% w/v sodium deoxycholate, 2 mM EDTA at pH 8.0), and 2x with TE. After the last wash, supernatant was removed and beads were resuspended in 1 mM Tris at pH 8.0 and RNAse A (0.1µg/µl) and incubated 30 min at 37°C. To reverse the cross-linking, beads were incubated overnight at 65°C in 100 mM Tris-HCl at pH 8.0, 20 mM EDTA at pH 8.0, 2% SDS, and proteinase K (0.5 μg/μL). The DNA was purified with SureClean (Bioline) and DNA concentration was measured with Qubit dsDNA HS Assay Kit.

Libraries were generate with 3ng of ChIPed DNA using NEBNext Ultra II DNA Library Prep Kit for Illumina (New England Biolabs), size was determined with Agilent Bioanalyzer DNA high sensitivity, and sequencing was performed on an Illumina NovaSeq 6000 instrument according to the manufacturer’s instructions. Read quality of ChIP-seq data was assessed using FastQC (v0.11.5) and reads were aligned to the human genome version 38 using Bowtie2 (v2.0.2). MACS2 (v2.1.2) was used to call broad peaks from BAM files.

**DNA extraction and bisulfite conversion**

Cell pellets were thawed on dry ice and homogenized with a 20G needle in Qiagen Buffer RLT Plus (QIAGEN Inc.) with β-mercaptoethanol. DNA extractions were performed with the Qiagen AllPrep DNA/RNA Mini Kit as per manufacturer’s instructions. DNA quantity and purity were assessed by spectrophotometry.

1.5 μg of DNA per sample was split into two 750 ng aliquots for sodium bisulfite (BS) conversion or oxidative bisulfite (oxBS) conversion using the NuGEN TrueMethyl oxBS Module (NuGEN Technologies Inc.). One aliquot per sample was treated with the oxidation protocol while the second aliquot underwent mock oxidation; all samples were then sodium bisulfite converted according to the manufacturer’s instructions.

**DNA methylation data pre-processing**

Filtering was performed to remove internal SNP control probes (59) and cross-hybridizing probes (43,254). The pfilter function in the wateRmelon package was additionally used to remove sites with a detection *p*-value > 0.05 in 1% of samples (7,685) and sites with a bead count < 3 in 5% of samples (1,626). This left 813,589 EPIC probes in the final dataset.

The dasen function in wateRmelon was used to normalize the oxBS and BS data separately. Batch effects from chip, position, and passage were removed using the ComBat function in the SVA package. Hydroxymethylation (hmC) values were calculated by subtracting the normalized oxBS beta values from the normalized BS beta values. A threshold for hmC detectability (3.6%) was calculated using the 95% quantile of negative hmC values generated after subtraction; all probes which had a mean hmC level below this threshold were discarded.

**Multi-omic integration**

Genomic regions were split into enhancers (+/- 5000bp from H3K4me1 ChIP-seq peaks), promoters (+/- 1000bp from TSS), and gene bodies (remaining regions). DNAm and DNAhm *p*-values were weighted across genomic regions according to their significance (using Stouffer’s method), creating a combined *p*-value for each region. *p*-values from all modifications were then used to create weighted gene-level scores. Weights were chosen in order to return modules likely to contain differentially expressed genes, with equal probability of contribution from differential DNAm and/or differential DNAhm (total weight of 0.25 for each methylation type). Enhancers and promoters were weighted slightly higher than gene bodies due to the higher likelihood of DNAm at these regions influencing gene expression. Scores were annotated to a REACTOME protein-protein interaction network, and a spin-glass algorithm was used to identify modules that had altered DNAm, DNAhm, and expression for each comparison.

**Pyrosequencing**

1μL of BS or oxBS-converted DNA per 30μL reaction was used to perform PCR with HotStarTaq DNA Polymerase (QIAGEN Inc.) for 45 cycles at an annealing temperature of 58°C. PCR samples were visualized on a 1% agarose gel post-amplification to ensure integrity.

Pyrosequencing assays were designed with PyroMark^®^ Assay Design 2.0 software (QIAGEN Inc.; Table S1). 10μl PCR reaction, 70μl binding buffer, and 12μl sequencing reaction mix per sample was prepared with PyroMark^®^ Gold Q96 Reagents according to the manufacturer’s instructions. Sequencing was performed with the PyroMark^®^ Q96 ID (QIAGEN Inc.).

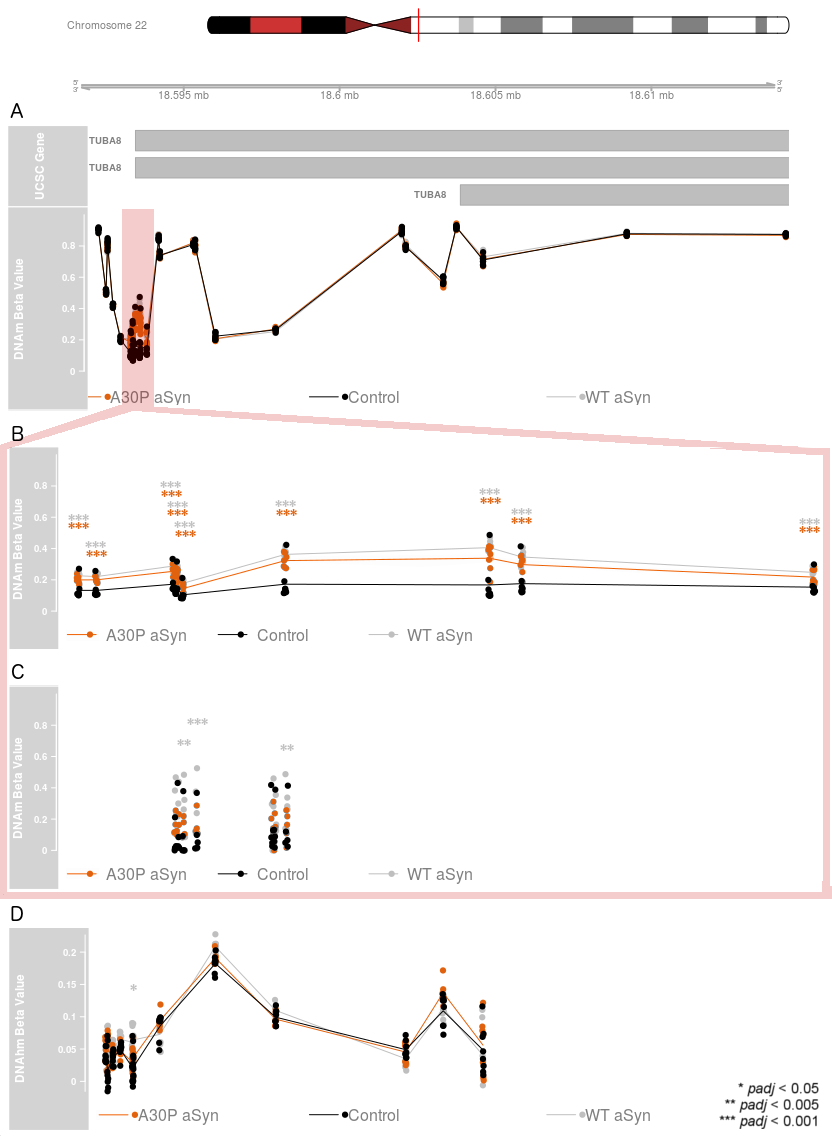

**Figure S1. *TUBA8* was differentially methylated in both LUHMES genotypes and differentially hydroxymethylated in WT aSyn cells.** Top: UCSC hg19 coordinates are shown, with *TUBA8* transcripts below. Bottom: Beta values are shown for each sample, coloured by genotype. Black: control cells; grey: WT aSyn cells; orange: A30P aSyn cells. (A) DNAm levels for all EPIC array probes across the *TUBA8* gene. (B) Differentially methylated region of *TUBA8*. (C) DNAm levels at five CpG sites in the *TUBA8* differentially methylated region measured by oxidative bisulfite pyrosequencing. (D) DNAhm levels for all EPIC array probes across the *TUBA8* gene. * *padj* < 0.05, ** *padj* < 0.005, *** *padj* < 0.001.

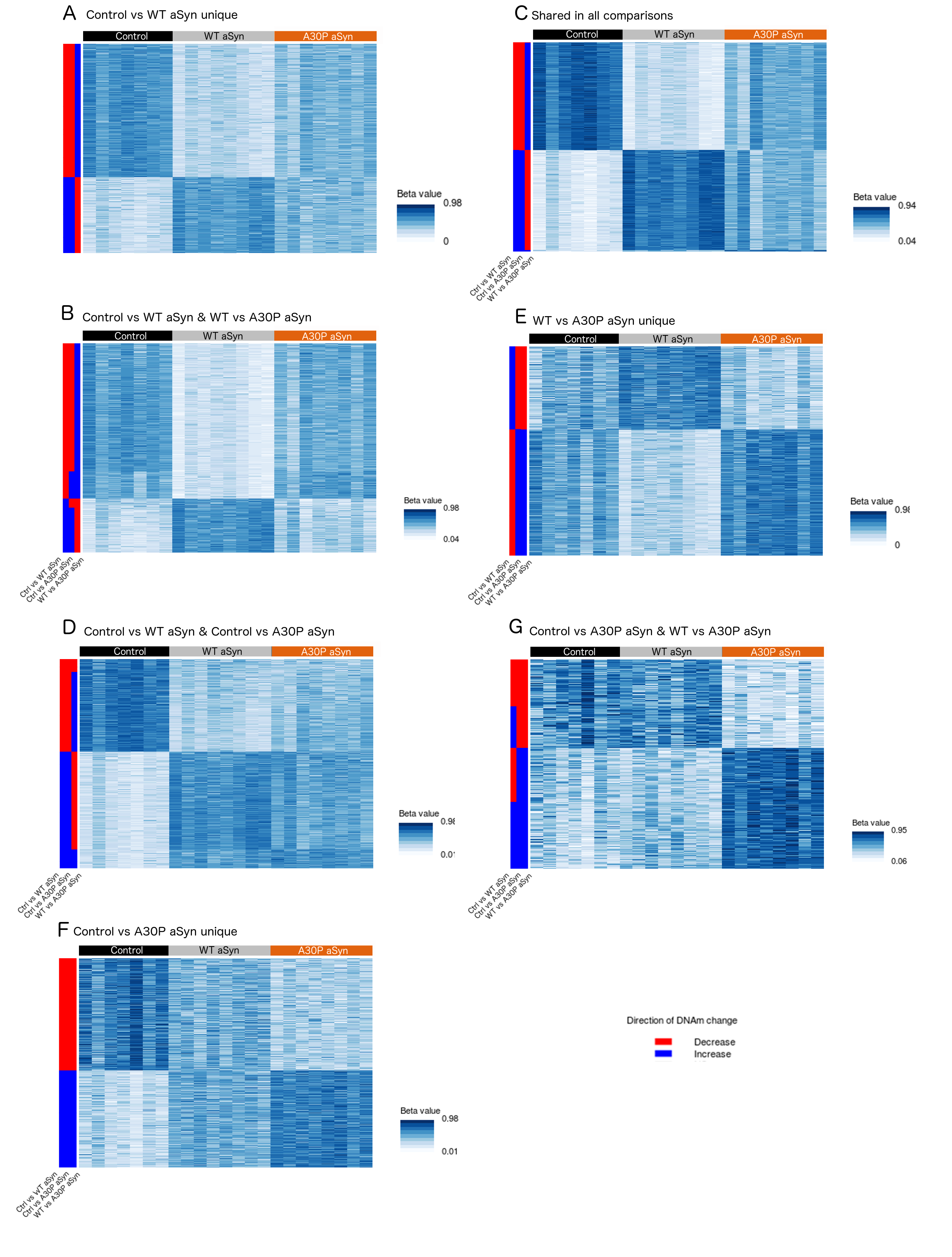

**Figure S2. DNAm levels of hits across comparisons.** Heat maps showing beta values for all probes differentially methylated in each comparison (|delta beta| >= 0.05 and *padj* <= 0.05). Row labels, left to right: Direction of DNAm change in control vs WT aSyn (decreased in WT aSyn: red, increased in WT aSyn: blue); direction of DNAm change in control vs A30P aSyn (decreased in A30P aSyn: red, increased in A30P aSyn: blue); direction of DNAm change in WT vs A30P aSyn (decreased in A30P aSyn: red, increased in A30P aSyn: blue). (A) 18,170 hits unique to control vs WT aSyn. (B) 3,791 hits shared only between control vs. WT and control vs. A30P aSyn. (C) 1,596 hits unique to control vs A30P aSyn. (D) 826 hits shared between all three comparisons. (E) 6,462 hits shared only between control vs. WT aSyn and WT vs. A30P aSyn. (F) 2,340 hits unique to WT vs A30P aSyn. (G) 232 hits shared only between control vs. A30P aSyn and WT vs. A30P aSyn.

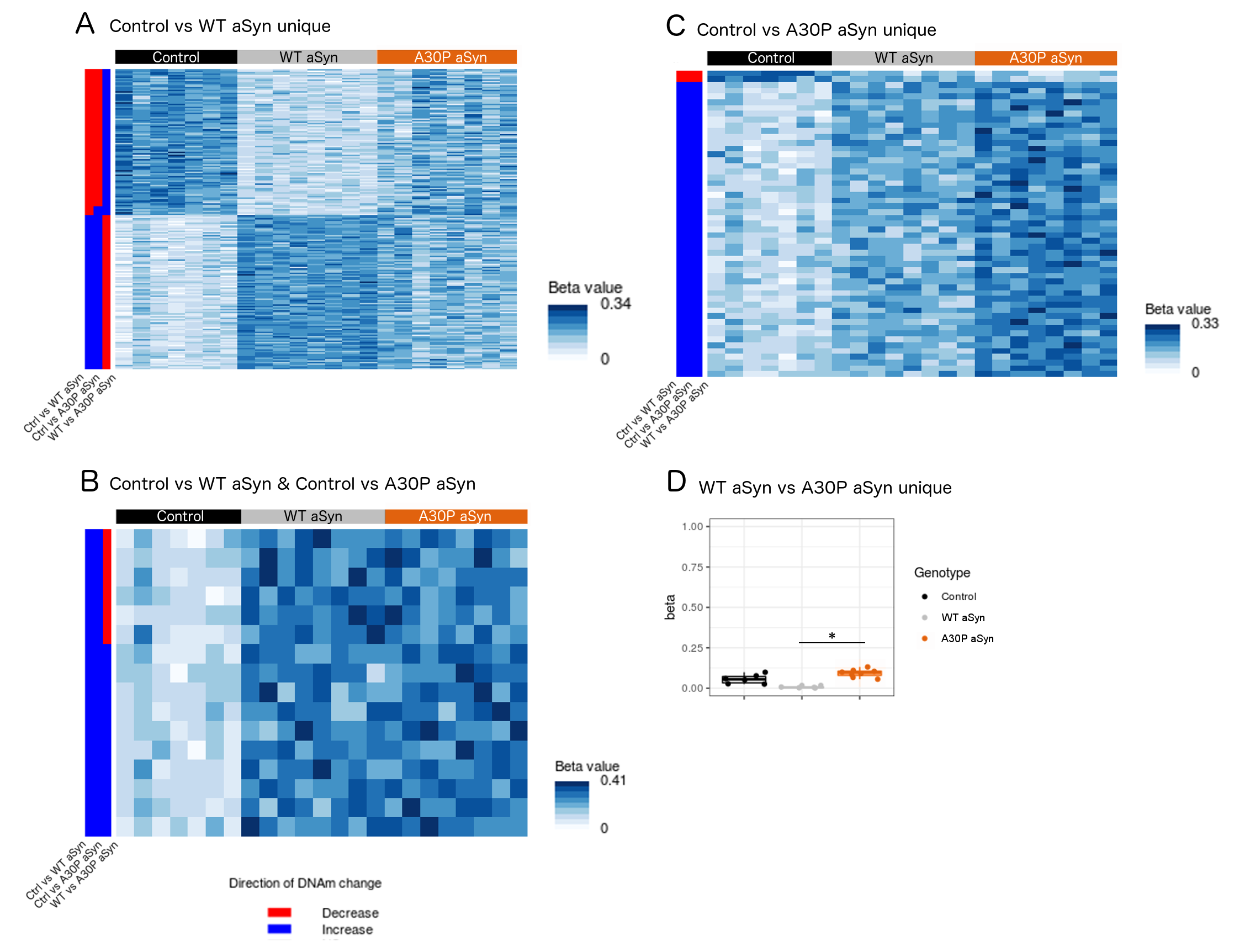

**Figure S3. DNAhm levels of hits across comparisons.** Heat maps showing beta values

for all probes differentially hydroxymethylated in each comparison (|delta beta| >= 0.05 and *padj*

<= 0.05). Row labels, left to right: Direction of DNAhm change in control vs WT aSyn (decreased in WT aSyn: red, increased in WT aSyn: blue); direction of DNAhm change in control vs A30P aSyn (decreased in A30P aSyn: red, increased in A30P aSyn: blue); direction of DNAhm change in WT vs A30P aSyn (decreased in A30P aSyn: red, increased in A30P aSyn: blue). (A) 199 hits unique to control vs WT aSyn. (B) 16 hits shared only between control vs. WT aSyn and control vs. A30P aSyn. (C) 51 hits unique to control vs A30P aSyn. (D) Box plots showing DNAhm levels for one CpG site unique to the WT vs A30P aSyn comparison. Black: control; grey: WT aSyn; orange: A30P aSyn. * *padj* < 0.05.

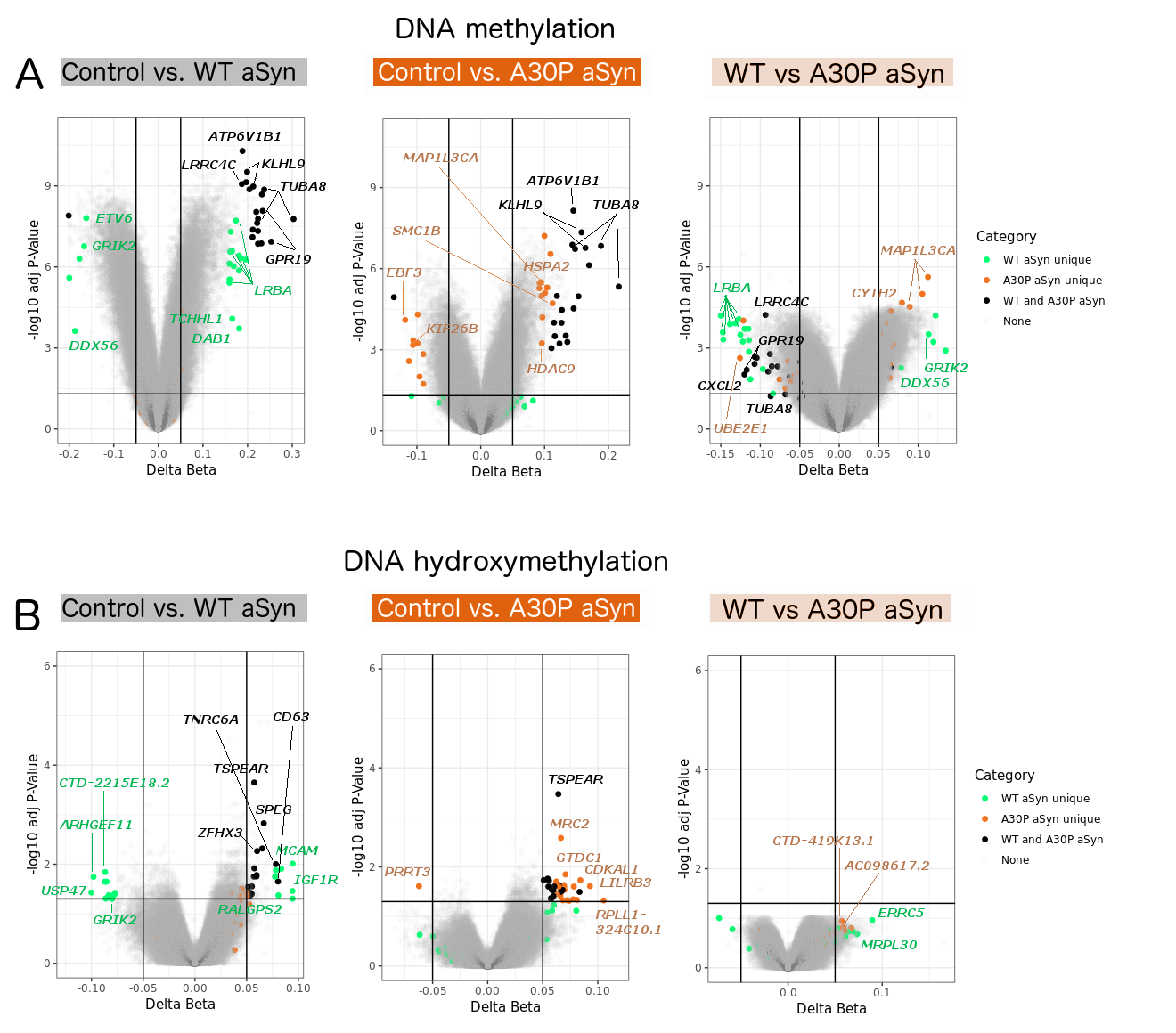

**Figure S4. Top probes unique to control vs WT aSyn, unique to control vs A30P aSyn, and shared between comparisons.** Probes were ranked by absolute delta beta. Green: top 20 probes unique to control vs WT aSyn comparison; orange: top 20 probes unique to control vs A30P aSyn comparison; black: top 20 probes shared between control vs WT aSyn and control vs A30P aSyn comparisons; grey: probes not included in above categories. (A) DNAm, (B) DNAhm.

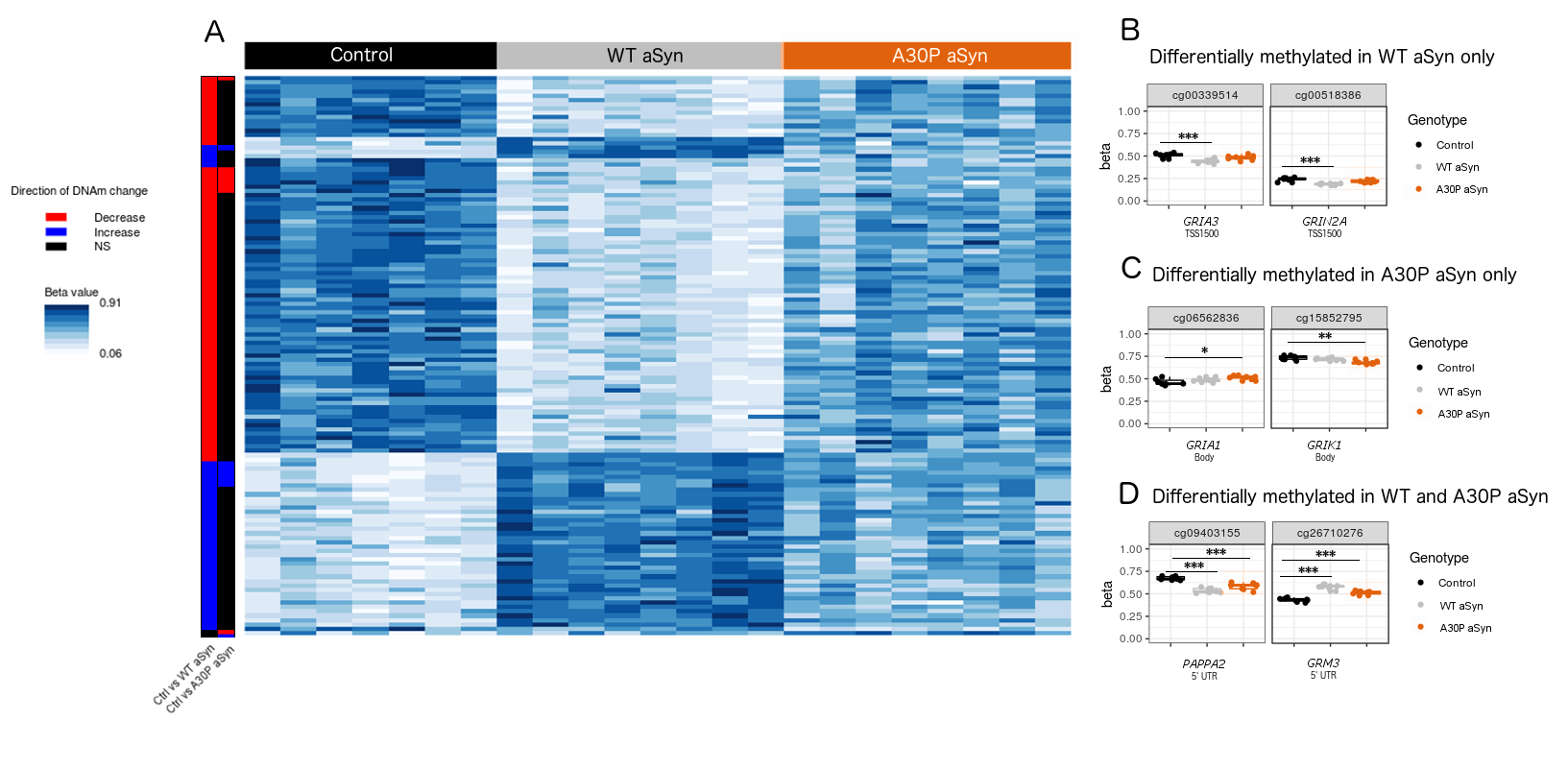

**Figure S5. Differentially methylated sites associated with glutamate receptor signaling genes in WT and A30P aSyn cells.** (A) Heat map showing beta values for all probes annotated to GO:0007215 (glutamate receptor signaling pathway) which were differentially methylated in either control vs. WT or control vs. A30P aSyn analyses. Row labels, left to right: Probes that passed significance thresholds (absolute delta beta >= 0.05 and *padj* <= 0.05) in control vs WT aSyn comparison (decreased DNAm in WT aSyn red; increased DNAm in WT aSyn blue; non-significant: black). Probes that passed significance thresholds in control vs A30P aSyn comparison (decreased DNAm in A30P aSyn: red; increased DNAm in A30P aSyn: blue; non-significant: black). (C) Representative examples of glutamate signaling-related CpG sites differentially methylated only in WT aSyn cells. (D) Representative examples of glutamate signaling-related CpG sites differentially methylated only in A30P aSyn cells. (E) Representative examples of glutamate signaling-related CpG sites differentially methylated in both genotypes. Black: control cells; grey: WT aSyn cells; orange: A30P aSyn cells. * *padj* < 0.05, ** *padj* < 0.005, *** *padj* < 0.001.

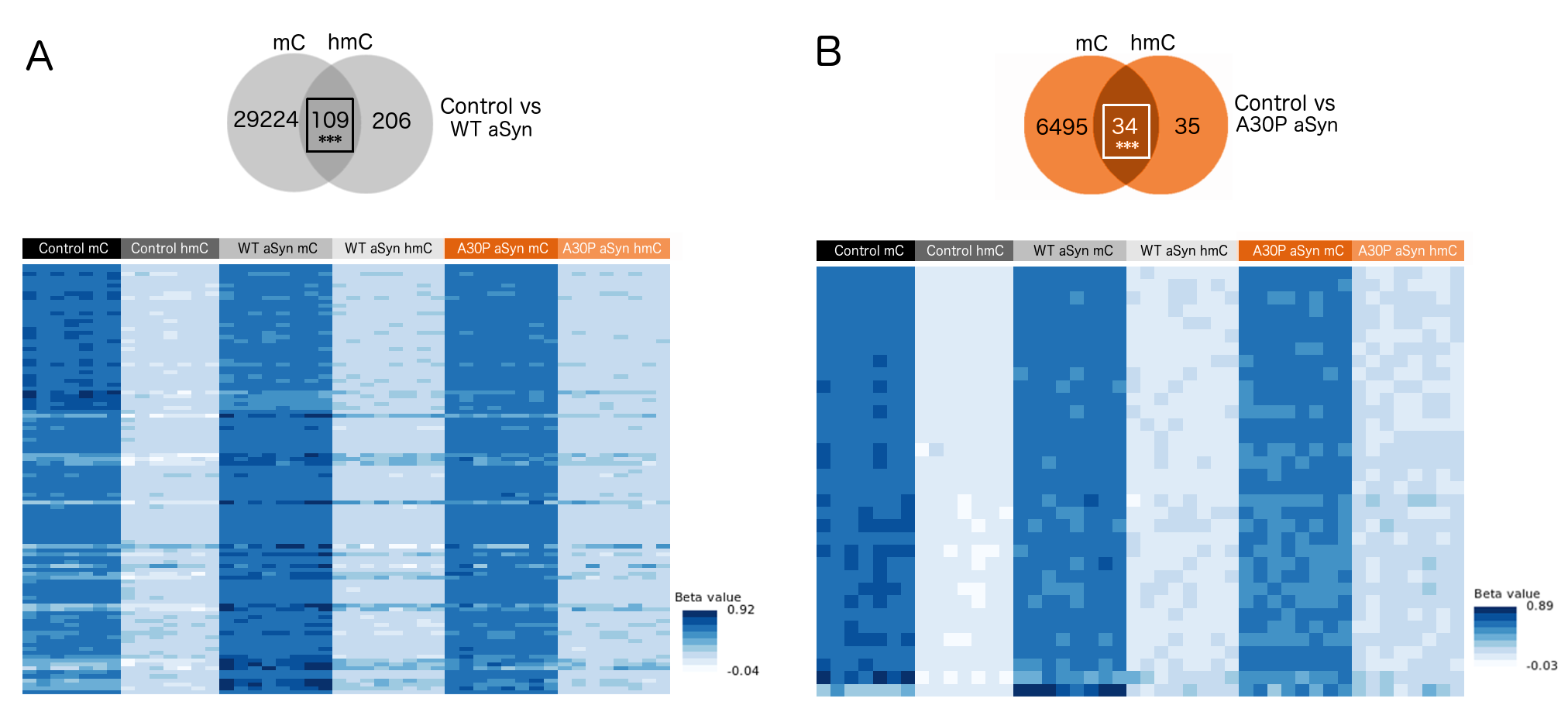

**Figure S6. DNAm and hydroxymethylation levels of hits in both modifications**. Heat maps

showing beta values for all probes differentially methylated and hydroxymethylated in each comparison

(|delta beta| >= 0.05 and *padj* <= 0.05). (A) DNAm and DNAhm levels of 109 hits with changes in both modifications in control vs WT aSyn. (B) DNAm and DNAhm levels of 34 hits with changes in both modifications in control vs A30P aSyn.

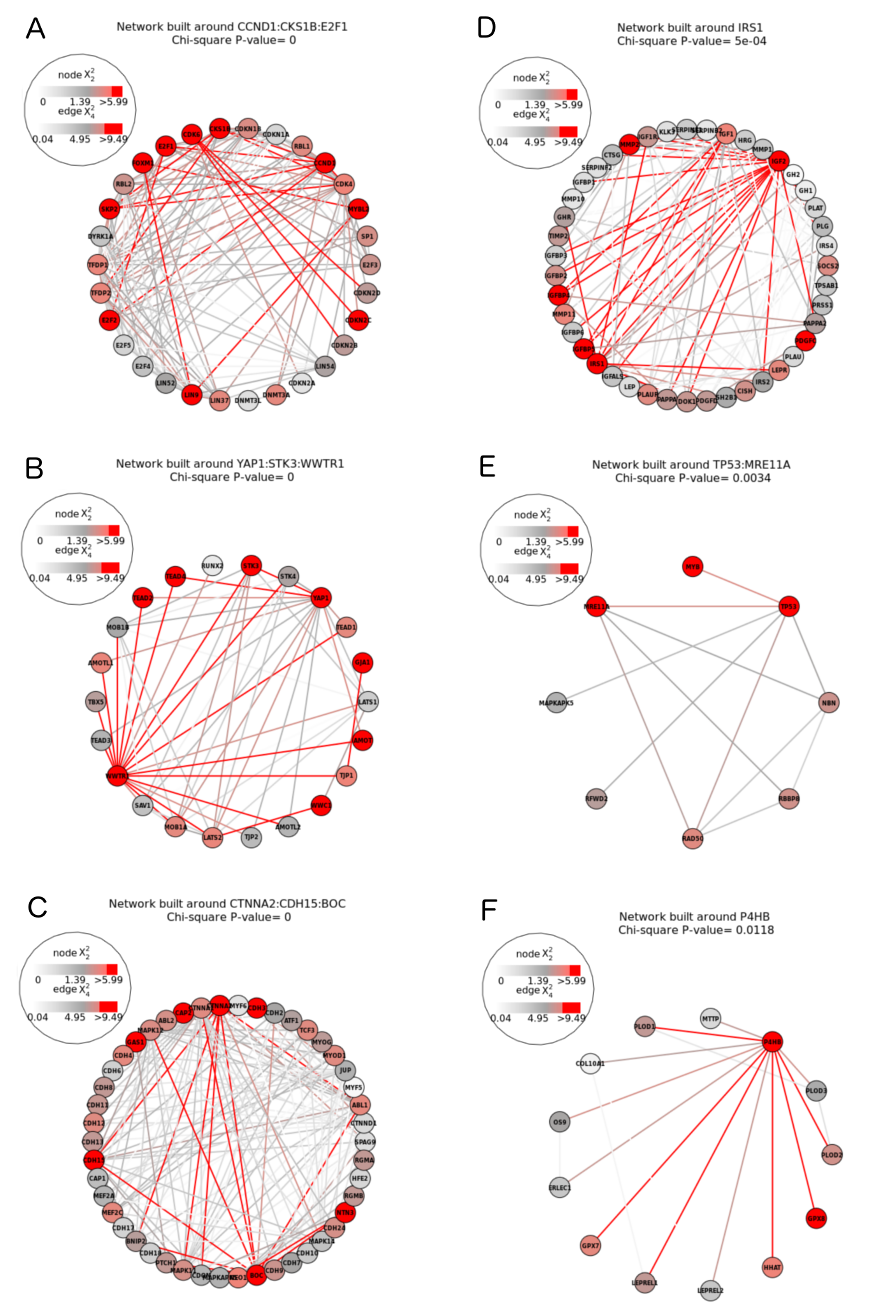

**Figure S7. SMITE modules dysregulated in WT aSyn cells.** Red fill: high scoring gene; red line: high scoring interaction (average score between genes). One representative module is shown for each of (A) cell cycle regulation, (B) apoptosis, (C) neuron development, (D) insulin signaling, (E) DNA damage repair, and (F) chaperone activity.

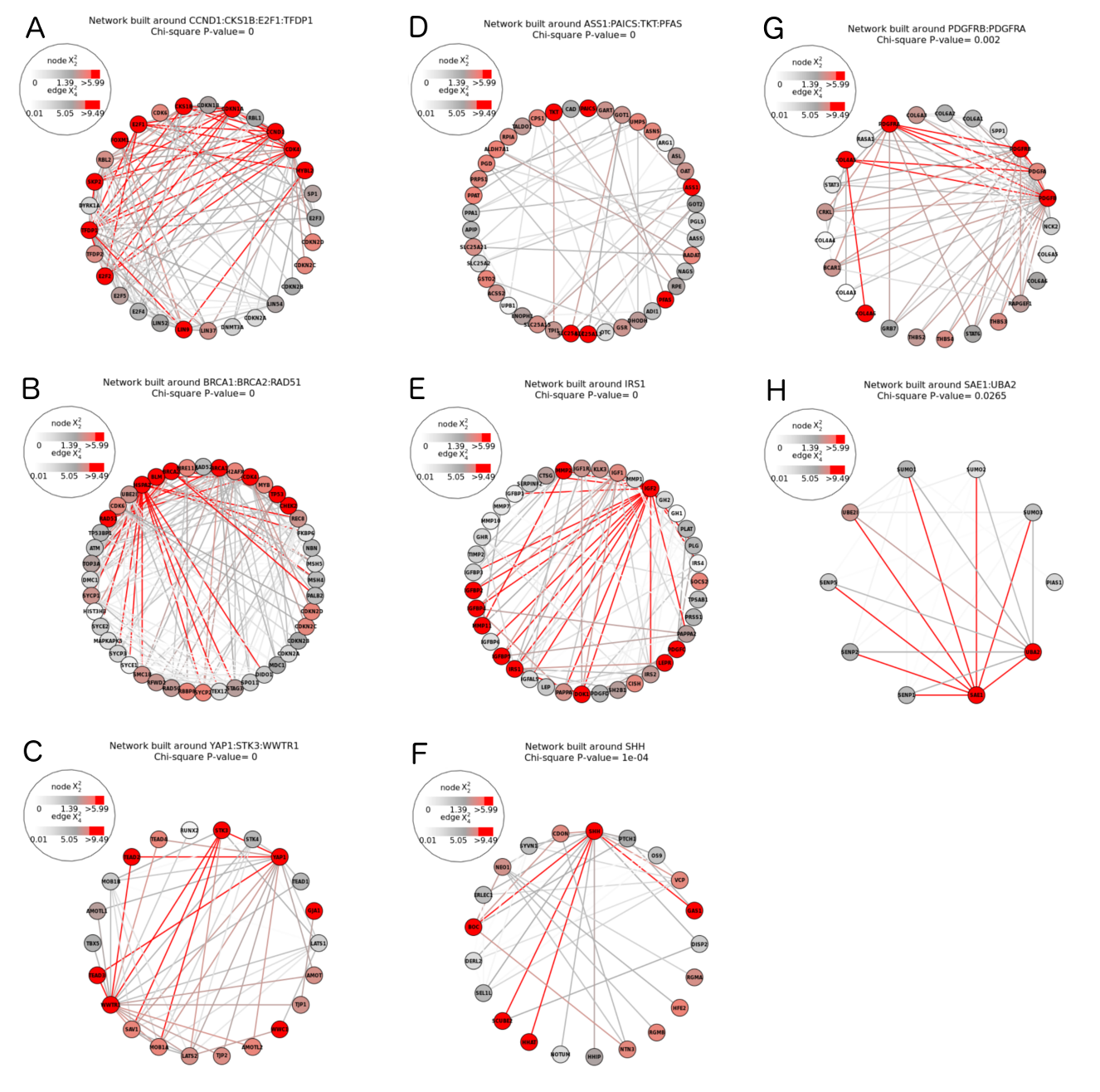

**Figure S8. SMITE modules dysregulated in A30P aSyn cells.** Red fill: high scoring gene; red line: high scoring interaction (average score between genes). One representative module is shown for each of (A) cell cycle regulation, (B) DNA damage repair, (C) apoptosis, (D) urea cycle, (E) insulin signaling, (F) neuron development, (G) PDGF signaling, and (H) sumoylation.

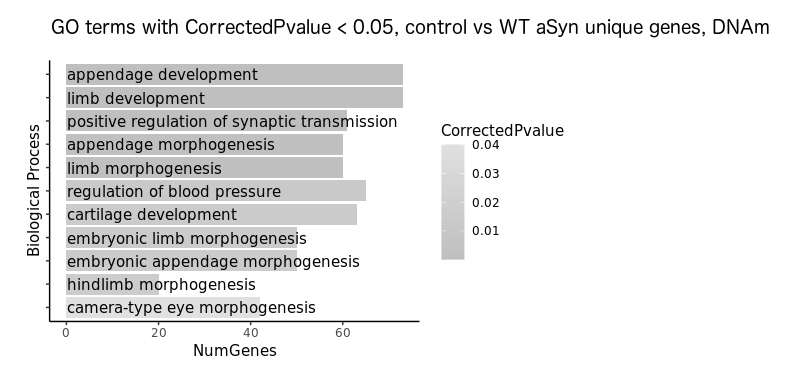

**Figure S9. GO biological process over-representation analysis for genes differentially methylated in control vs WT aSyn analysis.** 11 GO biological process terms signficant at corrected p-value < 0.05 are shown. Dark fill: more significant; light fill: less significant. NumGenes: number of genes in gene set.
